# Separable Fully Connected Layers Improve Deep Learning Models For Genomics

**DOI:** 10.1101/146431

**Authors:** Amr Mohamed Alexandari, Avanti Shrikumar, Anshul Kundaje

**Affiliations:** Departments of Computer Science and Genetics, Stanford University

## Abstract

Convolutional neural networks are rapidly gaining popularity in regulatory genomics. Typically, these networks have a stack of convolutional and pooling layers, followed by one or more fully connected layers. In genomics, the same positional patterns are often present across multiple convolutional channels. Therefore, in current state-of-the-art networks, there exists significant redundancy in the representations learned by standard fully connected layers. We present a new separable fully connected layer that learns a weights tensor that is the outer product of positional weights and cross-channel weights, thereby allowing the same positional patterns to be applied across multiple convolutional channels. Decomposing positional and cross-channel weights further enables us to readily impose biologically-inspired constraints on positional weights, such as symmetry. We also propose a novel regularizer and constraint that act on curvature in the positional weights. Using experiments on simulated and *in vivo* datasets, we show that networks that incorporate our separable fully connected layer outperform conventional models with analogous architectures and the same number of parameters. Additionally, our networks are more robust to hyperparameter tuning, have more informative gradients, and produce importance scores that are more consistent with known biology than conventional deep neural networks.

**Availability:** Implementation: https://github.com/kundajelab/keras/tree/keras_1

A gist illustrating model setup is at: goo.gl/gYooaa

## 1 INTRODUCTION

Convolutional neural networks are being successfully applied to predict important regulatory patterns from DNA sequences (Alipanahi et al. [2015], Kelley et al. [2016]). In such networks, a fully connected layer is connected to all the activations produced by the stack of convolutional layers. The job of the fully connected layer is to learn to combine these activations in a way that allows it to make a meaningful prediction about the input. In genomics, the same positional activation patterns are relevant across multiple convolutional channels, and different neurons in the fully connected layer must learn these patterns independently (Shrikumar et al. [2017b]). We develop separable fully connected (SFC) layers to make better use of the parameters in the network and enable biologically valid learning, in which genomic priors can be readily incorporated into the models.

## 2 METHODS

### 2.1 SEPARABLE FULLY-CONNECTED LAYERS

In deep learning models that have 1-d convolutions, the output of a stack of convolutional layers is typically reshaped from a 3-d tensor with dimensions (samples, length, channels) into a 2-d tensor with dimensions (samples, length × channels), and is supplied to a fully connected (FC) layer with a 2-d weights matrix of dimensions (num_outputs, length × channels). The FC layer takes the matrix product of the input tensor with its weights matrix to get a tensor with dimensions (samples, num_outputs). As illustrated in Figure 1, there are two major drawbacks to this approach. The first is cross-channel redundancy; the same positional patterns are often shared across multiple convolutional channels, so the FC layer must relearn these patterns for each channel. The second is redundancy between different neurons; every neuron in the FC layer must relearn all positional patterns independently.

**Figure 1:**
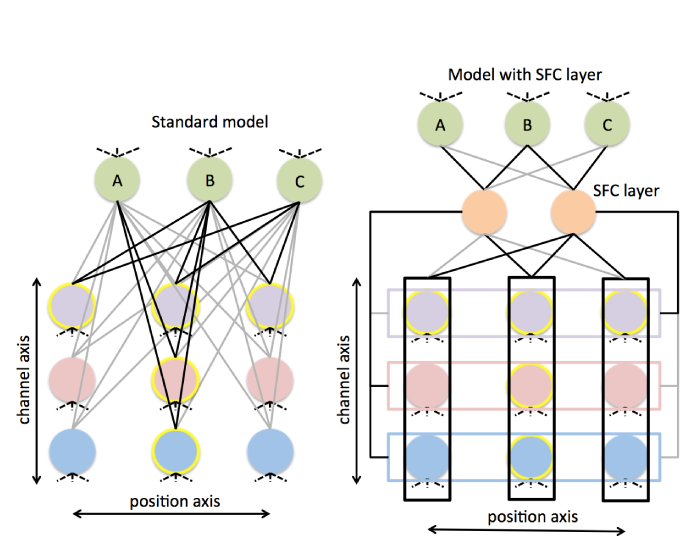
SFC layer overview. Toy example with 3 convolutional channels (blue, pink, and purple) corresponding to 3 different TFs. Pink and blue TFs bind in center of region, purple TF binds anywhere. Yellow halos indicate which positions are important for which TFs. Black edges correspond to weights of 1, grey edges correspond to a weights of 0. Neuron A in the FC layer responds to pink and blue TFs, neuron B responds to all TFs, neuron C responds only to purple TF. 2 kinds of redundancy exist: (1) pink and blue TFs have similar positional binding patterns, but these are learned for each channel separately, and (2) neurons A & B and B & C have subsets of TFs in common, but every neuron learns all weights independently. Standard FC layer has 9*3 = 27 parameters. Network with the SFC layer represents equivalent function with 3*2+ 6*2 = 18 parameters.

We address this problem by inserting a layer called a separable fully connected (SFC) layer between the convolutional layers and the existing FC layer. The SFC layer learns two separate two-dimensional tensors: *W*_*pos*_ with dimensions (num_outputs, length), and *W*_*chan*_ with dimensions (num_outputs, channels). It takes the outer product of *W*_*pos*_ and *W*_*chan*_ to form a 3-d tensor with dimensions (num_outputs, length, channels), and collapses the last two dimensions of this tensor to get a 2-d weights matrix with dimensions (num_outputs, length channels), called *W*_*out*_. The output is obtained by taking the matrix product of *W*_*out*_ with the input tensor of dimensions (samples, length *×* channels) as is done in standard FC layers.

We initialize *W*_*pos*_ and *W*_*chan*_ such that the variance of entries of *W*_*out*_ matches a glorot_uniform initialization (Glorot and Bengio [2010]) of a matrix with the same dimensions. Specifically, glorot_uniform would initialize the entries of *W*_*out*_ to be in the range 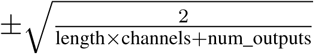so we initialize entries of *W*_*pos*_ and *W*_*chan*_ to be in the range 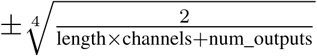. We found that correct initialization was critical for good performance.

It is important to note that the SFC layer is not followed by a nonlinearity; rather it feeds directly into the existing FC layer. Thus, any function that can be represented by the architecture containing the SFC layer can also be represented by the architecture lacking the SFC layer, because a linear function of a linear function is still linear. Therefore, any improvements are exclusively due to the efficient parameter usage that results from separating *W*_*pos*_ and *W*_*chan*_.

The change in the number of parameters that comes from introducing a SFC layer can be computed as follows: in the absence of a SFC layer, a FC layer that comes after a stack of convolutional layers has num_outputs × (length × channels) parameters. If we introduce a SFC layer with *n* neurons before the FC layer, the SFC layer will have *n*(length + channels) parameters, and the FC layer will have (num_outputs × *n*) parameters, for a total of *n*(length + channels + num_outputs) parameters. We can then solve for *n* to keep the number of parameters in the new model the same.

### 2.2 SYMMETRY AND CURVATURE REGULARIZATION/CONSTRAINT

Separating *W*_*pos*_ and *W*_*chan*_ in the SFC layer allows us to impose biologically-inspired constraints and penalties on *W*_*pos*_. For example, we can constrain *W*_*pos*_ to be symmetric around the center, which we would expect to be appropriate for tasks where the input regions are not oriented according to any directional property. We can also impose curvature regularization (CR) on *W*_*pos*_ by applying an *ℓ*_1_ penalty to the second differences of adjacent columns in *W*_*pos*_, where first difference (*d*_1_) is defined as *d*_1_[*i*] = *W* [*i*] − *W* [*i* − 1] ∀*i* and second difference (*d*_2_) is defined as *d*_2_[*i*] = *d*_1_[*i*] − *d*_1_[*i* − 1] ∀*i*. Analogously, we can apply a curvature constraint (CC) by enforcing -*α ≤ d*_2_[*i*] ≤ *α* for some positive constant *α*. If *W*_*pos*_ is symmetric, we expect a change in curvature around the center. To avoid penalizing this expected change in curvature, we apply our regularization/constraint in conjunction with the symmetry constraint, which allows us to learn and regularize/constrain only half of *W*_*pos*_ and reflect it to obtain the other half.

### 2.3 DATASETS

For the simulated transcription factor (TF) binding dataset, we use the same simulated (balanced) dataset from Shrikumar et al. [2017a], and train on 6,400 examples. For *in vivo* TF binding data, we use chromatin immunoprecipitation sequencing (ChIP-seq) datasets for Gm12878 obtained from the ENCODE Consortium [2012] as described with the relevant labels and data imbalance in Shrikumar et al. [2017b].

For accessibility datasets, the output consists of a binary vector where each entry indicates accessibility in a particular cell type. For the simulated accessibility dataset, we use DNase peaks from the ENCODE Consortium [2012] for two cell types: Lymphoblastoids data (indexed E116) and Leukemia cell data (indexed E123). The dataset is generated by (1) merging the DNase peaks from both cell types (2) padding each merged region with 100bp on either side (3) sliding windows of size 200bp tiling the padded merged region, with a stride of 50bp(4) intersecting each 200bp segment with the original DNase files from each cell type; regions that have 50% overlap are labeled with a 1, otherwise with a 0 (5) expanding the 200bp segments to be 1000bp (6) identifying the summits of TF ChIP-seq peaks from ENCODE, intersecting these summits with the 1000bp regions produced by (5) to identify putative locations of motifs (7) for each 1000bp DNase region, dinucleotide shuffling the underlying DNA sequence and inserting motifs associated with each TF at the location of the summit. We train on 2,178,417 examples and the ratios negatives:positives are 1.32 and 1.38 for E116 and E123 respectively. For the *in vivo* accessibility dataset, we use a hematopoiesis dataset measuring accessibility in 16 cell types from Corces et al. [2016], and train on 837,977 examples. The ratios negatives:positives in the 16 tasks range from 2.00 to 13.69 with half the ratios between 2 and 3.

### 2.4 MODEL ARCHITECTURE AND TRAINING

We trained all models using Keras (Chollet [2017]) with the Tensorflow backend using 1d convolutions. On all datasets we compare a standard model, a model with a SFC layer added before the FC layer, and a model with a symmetric SFC layer. The number of parameters in the model with a SFC layer is always less than or equal to the number in the corresponding model without a SFC layer. For both simulated and *in vivo* accessibility tasks, we use the Basset architecture, discussed by Kelley et al. [2016], as the standard model. When a SFC layer was added to this model, we used 1000 neurons. For the simulated TF binding dataset, the standard model had 2 convolutional layers (with ReLU) with 50 filters and filter width 11 each, followed by a max pooling layer with pooling length 10 and stride 5, followed by a FC layer with 50 neurons, followed by dropout (with probability 0.5), followed by an output layer with 3 sigmoid neurons. When we add a SFC layer to this model, we use 648 neurons. For the *in vivo* TF binding tasks, the standard model is described in section 3.4.2 in Shrikumar et al. [2017b]. When we add a separable fully connected layer to this model, we use 18 neurons. Models were trained using the Adam optimizer (Kingma and Ba [2014]) and a binary cross-entropy loss. The learning rate for Adam was 0.001 for all models except the standard model trained on simulated accessibility data, for which a learning rate of 0.0001 was used as it failed to learn with a learning rate of 0.001.

## 3 RESULTS

### 3.1 SFC LAYERS IMPROVE PERFORMANCE

We compared the performance of models with and without SFC layers on simulated and *in vivo* TF-binding datasets, as well as simulated and *in vivo* accessibility datasets. As illustrated in Figure 2, the median auROC/auPRC is consistently higher for models with a SFC layer than for models without. SFC layers with a symmetry constraint perform comparable to or better than those without, particularly on the simulated TF binding dataset and hematopoiesis dataset. One possible explanation is that the symmetry constraint is most helpful when the amount of training data is low. We applied CR/CC as described in Sec. 2.2 to models trained on the in-vivo hematopoiesis dataset and observed performance improvements over applying *ℓ*_1_ regularization on the first differences of *W*_*pos*_. We also noted that models with a SFC layer had more informative gradient updates and thus trained in fewer epochs, as shown in Figure 3.

**Figure 3:**
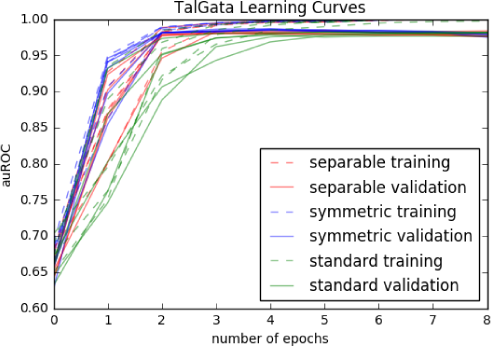
Models with SFC layer learn in fewer epochs. Learning curves show auROC (averaged across all outputs) for simulated TF-binding dataset for standard model, model with SFC, and model with symmetric SFC.

**Figure 2:**
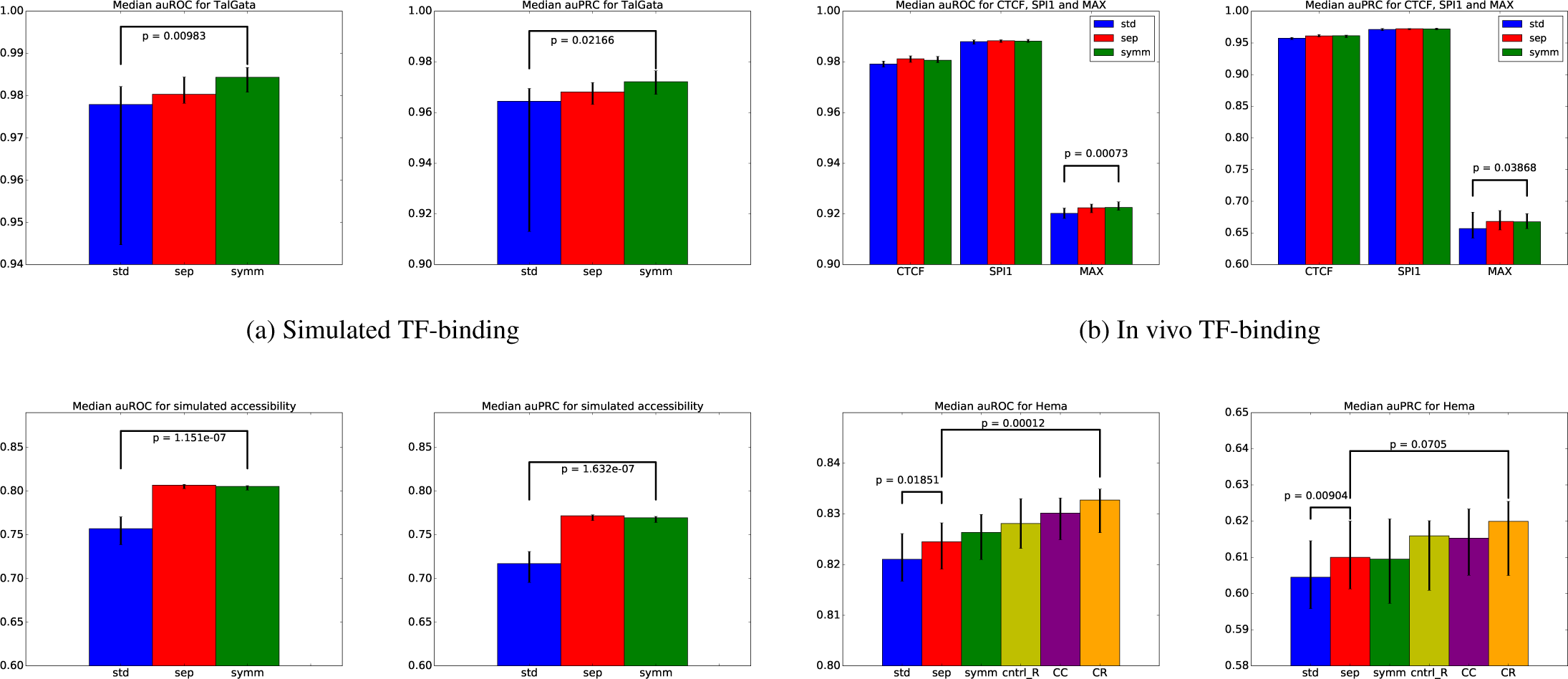
Separable fully-connected layers improve performance. The performance of different models was evaluated on different prediction tasks (see **Sec. 2.4**). Median auROC/auPRC (averaged across all outputs) on the validation set was computed over 10 random seeds; black error bars indicate min and max performance over different random seeds. `std`=standard model, `sep`=model with SFC layer, `symm`=model with SFC layer and symmetry constraint. First difference regularization and curvature regularization/constraint (CR/CC; **Sec. 2.2**) were applied to models trained on the hematopoiesis accessibility dataset containing a symmetric SFC layer; ctrl_R = first difference regularization of 10.0, CC = CC of 0.0001, and `CR` = CR of 10.0. CR of 1.0 and 100.0 performed worse than 10.0, CC of 0.001 and 0.00001 performed worse than 0.0001, and first difference regularization of 1.0 and 100.0 performed worse than 10.0. p-values were computed using 1-sided paired t-tests with a pairing over each random seed.

On both the simulated TF-binding dataset and simulated accessibility dataset, we also observed that the stability of performance across different random seeds was far better for models with a SFC layer (black error bars indicate minimum and maximum performance across 10 random seeds). In fact, models without a SFC failed to train (i.e. obtained 0.5 auROC) on the simulated accessibility dataset when the default learning rate (0.001) was used for the Adam optimizer. Models with a SFC layer successfully learned under both learning rates (0.0001 and 0.001).

### 3.2 SFC LAYERS GIVE MORE BIOLOGICALLY CORRECT IMPORTANCE SCORES

We visualized the mean importance at each position in the sequence for the best-performing model trained on the simulated TF-binding dataset in Figure 4a. For the simulated dataset, motifs had been embedded uniformly within the sequence; thus, an ideal model would give equal importance to all positions in the sequence. The models with separable FC layers come closest to this, particularly when using a symmetry constraint. We also visualized per-position importance on a model trained on the in-vivo CTCF dataset (Figure 4b). Despite the large training set size, importance scores for the standard model without a SFC layer are not perfectly smooth or symmetric.

**Figure 4:**
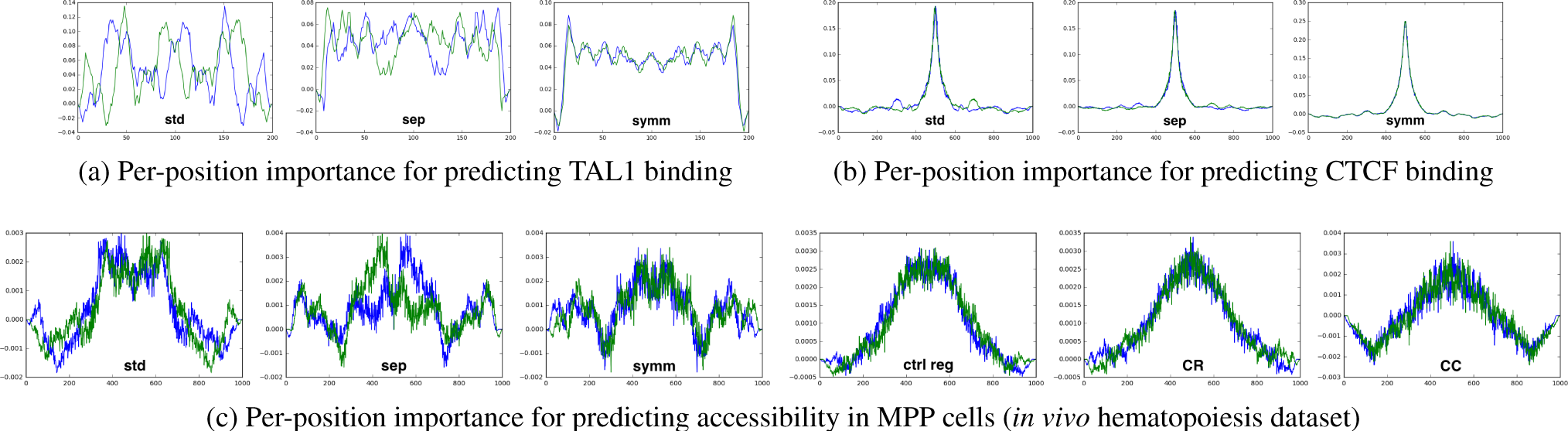
SFC layers improve per-position importance scores. Importance scores were computed on the input sequences using gradient × input as described in Shrikumar et al. [2017a] for the best performing model over 10 random seeds. Scores at each position were averaged across multiple sequences and visualized. Blue lines represent the original per-position importance. Green lines are mirror images of the corresponding blue lines (they are included to aid with visually assessing symmetry). **a**: per-position scores for TAL1 task in simulated TF-binding, averaged over 100K generated sequences. **b**: per-position scores for in-vivo CTCF task, averaged over regions in the testing set. **c**: per-position scores for MPP accessibility task in hematopoiesis dataset, averaged over 10K regions.

We performed a similar visualization of per-position importance scores for models trained on the hematopoiesis dataset (Figure 4c). Note that CR/CC markedly improved the smoothness of the scores. As corroborating evidence, we visualized the weights of *W*_*pos*_ for the model with CC as well as for models with CR at increasing levels of CR (Figure 5). Models that used higher CR or the CC learned smoother positional weights.

**Figure 5:**
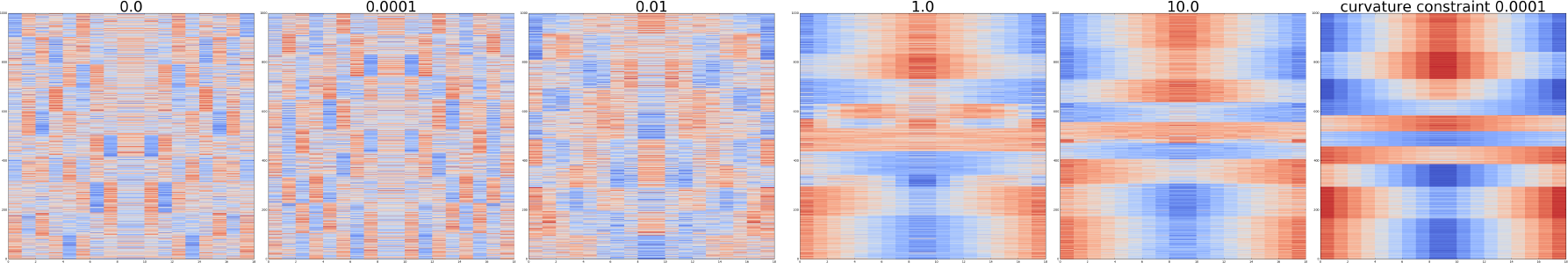
Curvature Regularization/Constraint produces smoother *W*_*pos*_. Weights of *W*_*pos*_ for models trained on the hematopoiesis dataset with different levels of CR are visualized. The last visualization is for CC. Each row corresponds to positional weights for a single output neuron. Strength of CR/CC is indicated on the top. Rows were mean-normalized to have unit magnitude, and clustered using k-means clustering with *k* = 10. Symmetry was enforced for all models.

## 4 CONCLUSION

We have demonstrated that standard fully connected layers are suboptimal due to redundancies in the learned patterns. By introducing separable fully connected layers into our models, we allow neurons to reuse positional patterns, thereby improving performance and learning cleaner representations. Finally, we show that imposing biologically meaningful constraints and penalties on the positional weights learned by the separable fully connected layers, such as symmetry and smoothness, further improves the networks.

## 5 ACKNOWLEDGMENTS

We thank Peyton Greenside for assistance with processing and learning models on the hematopoiesis dataset, and Irene Kaplow for feedback on the manuscript.

## 6 AUTHOR CONTRIBUTIONS

AMA implemented the layer and conducted all experiments under the mentorship of AS. AS & AMA worked out the concrete implementation details. AS & AK conceptualized the high-level idea for separable fully connected layers. AS conceived of the idea of applying curvature regularization and AMA conceived of the curvature constraint. AMA, AS, and AK wrote the manuscript.

## 7 FUNDING

AS is supported by a Howard Hughes Medical Institute International Student Research Fellowship and a Bio-X Bowes Fellowship. AK is supported by NIH grants DP2-GM-123485 and 1R01ES025009-02.

